# Spc110 N-Terminal Domains Act Independently to Mediate Stable γ-Tubulin Small Complex Binding and γ-Tubulin Ring Complex Assembly

**DOI:** 10.1101/311027

**Authors:** Andrew Lyon, Alex Zelter, Shruthi Viswanath, Alison Maxwell, Richard Johnson, King Clyde B. Yabut, Michael MacCoss, Trisha N. Davis, Eric Muller, Andrej Sali, David A. Agard

## Abstract

Microtubule (MT) nucleation *in vivo* is regulated by the γ-tubulin ring complex (γTuRC), an approximately 2-megadalton complex conserved from yeast to humans. In *Saccharomyces cerevisiae*, γTuRC assembly is a key point of regulation over the MT cytoskeleton. Budding yeast γTuRC is composed of seven γ-tubulin small complex (γTuSC) subassemblies which associate helically to form a template from which microtubules grow. This assembly process requires higher-order oligomers of the coiled-coil protein Spc110 to bind multiple γTuSCs, thereby stabilizing the otherwise low-affinity interface between γTuSCs. While Spc110 oligomerization is critical, its N-terminal domain (NTD) also plays a role that is poorly understood both functionally and structurally. In this work, we sought a mechanistic understanding of Spc110 NTD using a combination of structural and biochemical analyses. Through crosslinking-mass spectrometry (XL-MS), we determined that a segment of Spc110 coiled-coil is a major point of contact with γTuSC. We determined the structure of this coiled-coil segment by X-ray crystallography and used it in combination with our XL-MS dataset to generate an integrative structural model of the γTuSC-Spc110 complex. This structural model, in combination with biochemical analyses of Spc110 heterodimers lacking one NTD, suggests that the two NTDs within an Spc110 dimer act independently, one stabilizing association between Spc110 and γTuSC and the other stabilizing the interface between adjacent γTuSCs.

## Introduction

Spc110 plays a dual role in the budding yeast *Saccharomyces cerevisiae*, both connecting the γ-tubulin small complex (γTuSC) with the nuclear face of the spindle pole body (SPB) (Kilmartin, et al., 1993; Knop & Schiebel, 1997; Knop & Schiebel, 1998) and regulating assembly of γTuSC subassemblies into the microtubule-nucleating γ-tubulin ring complex (γTuRC) (Kollman, et al., 2010; Lin, et al., 2015). Our previous work has demonstrated that γTuSC self assembles via relatively low-affinity interactions which must be cooperatively stabilized by higher-order oligomers of Spc110 for efficient γTuRC assembly to occur both *in vitro* and *in vivo* (Lyon, et al., 2016). This higher-order assembly is thought to occur *via* coiled-coil mediated oligomerization, which is expected to be favorable *in vivo* due to high local concentrations of Spc110 at the SPB.

While higher-order oligomerization is clearly a crucial determinant of Spc110-mediated γTuRC stabilization, several observations indicated the importance of the 111-residue N-terminal domain (hereafter NTD^1-111^). If higher-order Spc110 oligomerization were the sole determinant of γTuRC assembly, Spc110 constructs that do not assemble beyond a dimer should not support γTuRC assembly. However, at high γTuSC concentration a dimeric Spc110 construct does induce assembly of γTuSCs into larger complexes, albeit with reduced average γTuSC number within the complexes and weaker affinity compared with tetrameric or larger Spc110 variants, implying an important role for other regions of Spc110 (Lyon, et al., 2016). Further, while the deletion of amino acid residues 1-34 of Spc110 leaves the protein functional *in vitro* and the mutant viable *in vivo*, a NTD^1-111^ deletion mutant is not viable *in vivo*, while supporting substantially reduced γTuRC assembly *in vitro*. Deletion of the NTD^1-111^ plus the centrosomin motif 1 (CM1^117-146^) abolishes γTuRC assembly *in vitro*. This interpretation is complicated by the presence of only a small portion of Spc110 in cryo-EM reconstructions of the disulfide-stabilized closed γTuSC filament, which suggests that the NTD^1-111^ fails to adopt a single stable conformation and hinders efforts to identify important interaction motifs (Kollman, et al., 2015; Greenberg, et al., 2016).

In this work, we refine the understanding of Spc110 in γTuRC assembly *via* structural and biochemical approaches. We use chemical crosslinking coupled with mass spectrometry (XL-MS) to define interactions between Spc110 and γTuSC. This allowed assignment of an Spc110 coiled-coil region (residues 164-203) as the density previously observed in a cryo-EM reconstruction (Kollman, et al., 2015). Integrative structure modeling indicates that data from cryo-EM, X-ray crystallography, and XL-MS is consistent with a single Spc110 NTD^1-111^ binding to γTuSC. Using our FRET assay for γTuRC assembly (Lyon, et al., 2016) and covalently linked Spc110 heterodimers, we present data indicating the two Spc110 NTD^1-111^s in the Spc110 dimer act independently to stabilize γTuRC assembly. One associates with the γTuSC to which the coiled coil region is bound, while the other stabilizes interactions with an adjacent laterally associated γTuSC.

## Results

### Defining Spc110-γTuSC interaction by XL-MS

The 6.9 Å cryo-EM reconstruction of Spc110-bound γTuSC filaments in a closed conformation revealed density consistent with two alpha-helices approximately 40 residues in length derived from Spc110, though the limited resolution prevents rigorous assignment of this density to any portion of Spc110. The Spc110^1-220^ construct used in the reconstruction contains a 45-residue high-probability coiled-coil segment at positions 164-208 (hereafter referred to as the N-terminal coiled-coil, or NCC^164-208^; see Figure S1), as well as a predicted helix within the centrosomin motif 1 (CM1^117-146^) (Figure 1A). Biochemical assays for γTuRC assembly indicate that the NTD^1-111^ of Spc110 contributes significantly to the stabilization of γTuRC. Cells expressing Spc110 lacking this domain are inviable (Lyon, et al., 2016). However, the cryo-EM map lacked any apparent density consistent with this region. There thus remains significant uncertainty about the interactions between these portions of Spc110 and γTuSC which we sought to resolve.

**Figure 1.**
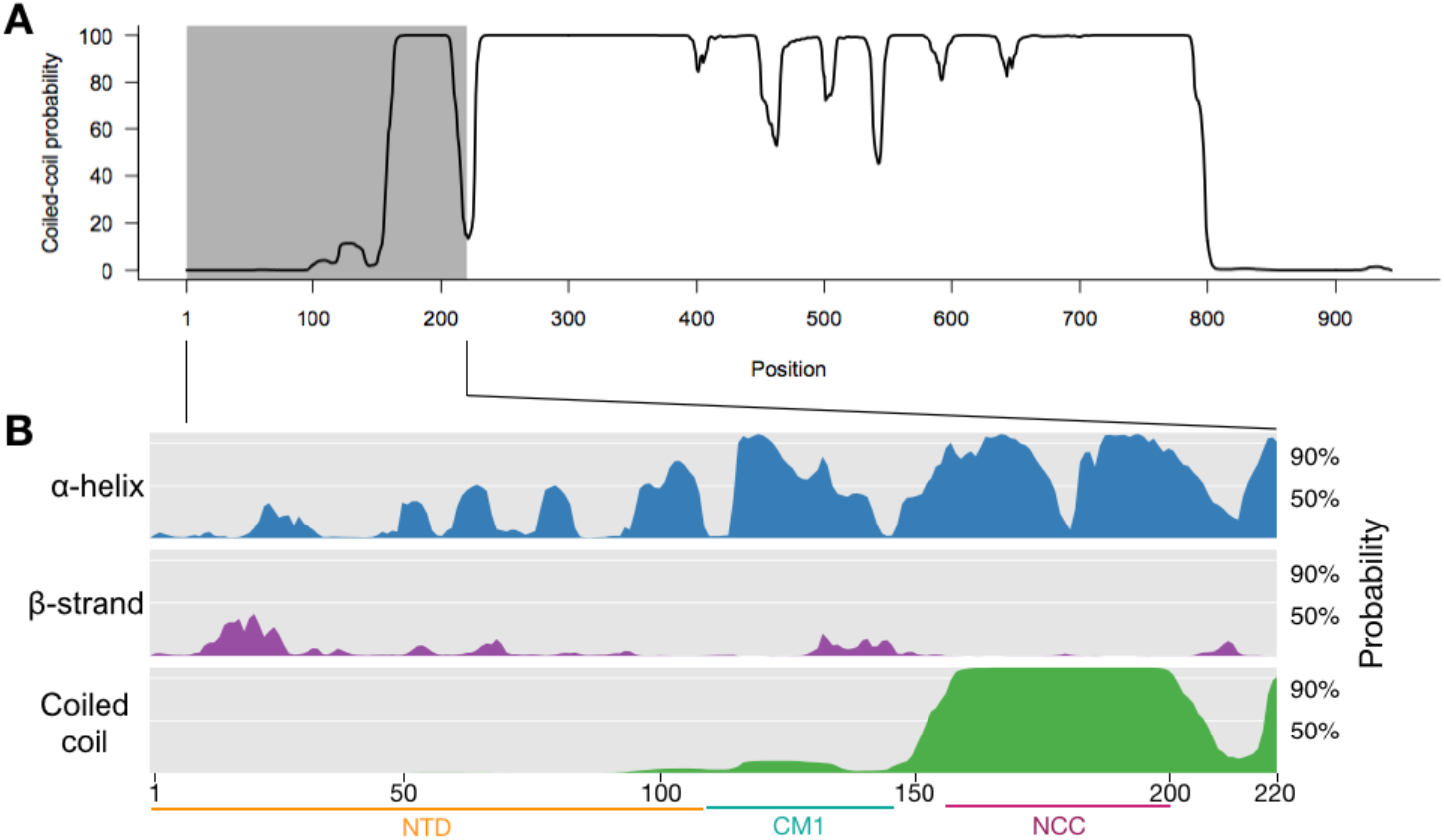
Spc110 domains and secondary structure. **A**. Spc110 coiled-coil prediction using MARCOIL. **B**. Spc110 N-terminal region secondary structure prediction, showing lack of predicted secondary structure for the first 111 residues. Also shown are CM1^117-146^ and the NCC^164-208^ regions.

To define these important interaction interfaces between Spc110 and γTuSC, we utilized chemical crosslinking coupled with mass spectrometry (XL-MS). We performed XL-MS using two Spc110 constructs. The Spc110^1-220^-GCN4 dimer construct stabilizes γTuRC assembly *in vitro* and *in vivo*, but relatively weakly compared with higher-order oligomers (Lyon, et al., 2016). The Spc110^1-401^-GST construct purified from baculovirus-infected insect cells forms large oligomers and stabilizes γTuRC assembly efficiently (Kollman et al. 2015). We used two chemical crosslinking reagents with different reactivities and linker lengths: disuccinimidyl suberate (DSS), a homo-bifunctional amine reactive reagent with an 11.4 Å aliphatic spacer, and 1-ethyl-3-(3-dimethylaminopropyl)carbodiimide hydrochloride (EDC), a so-called “zero-length” amine-carboxyl crosslinker. We identified a large number of high confidence crosslinked peptides in both Spc110 constructs with each crosslinking reagent.

We first focused on crosslinks between the N-terminal portions of Spc97 and Spc98 and Spc110 that would inform on the identity of the Spc110 alpha-helical densities observed in the cryo-EM map of Spc110-bound γTuSC. We observed a series of EDC crosslinks between the NCC^164-208^ domain and the N-terminal portions of Spc97 consistent with the coiled-coil-γTuSC interaction apparent in the cryo-EM map (Figure 2, red asterisks).

**Figure 2.**
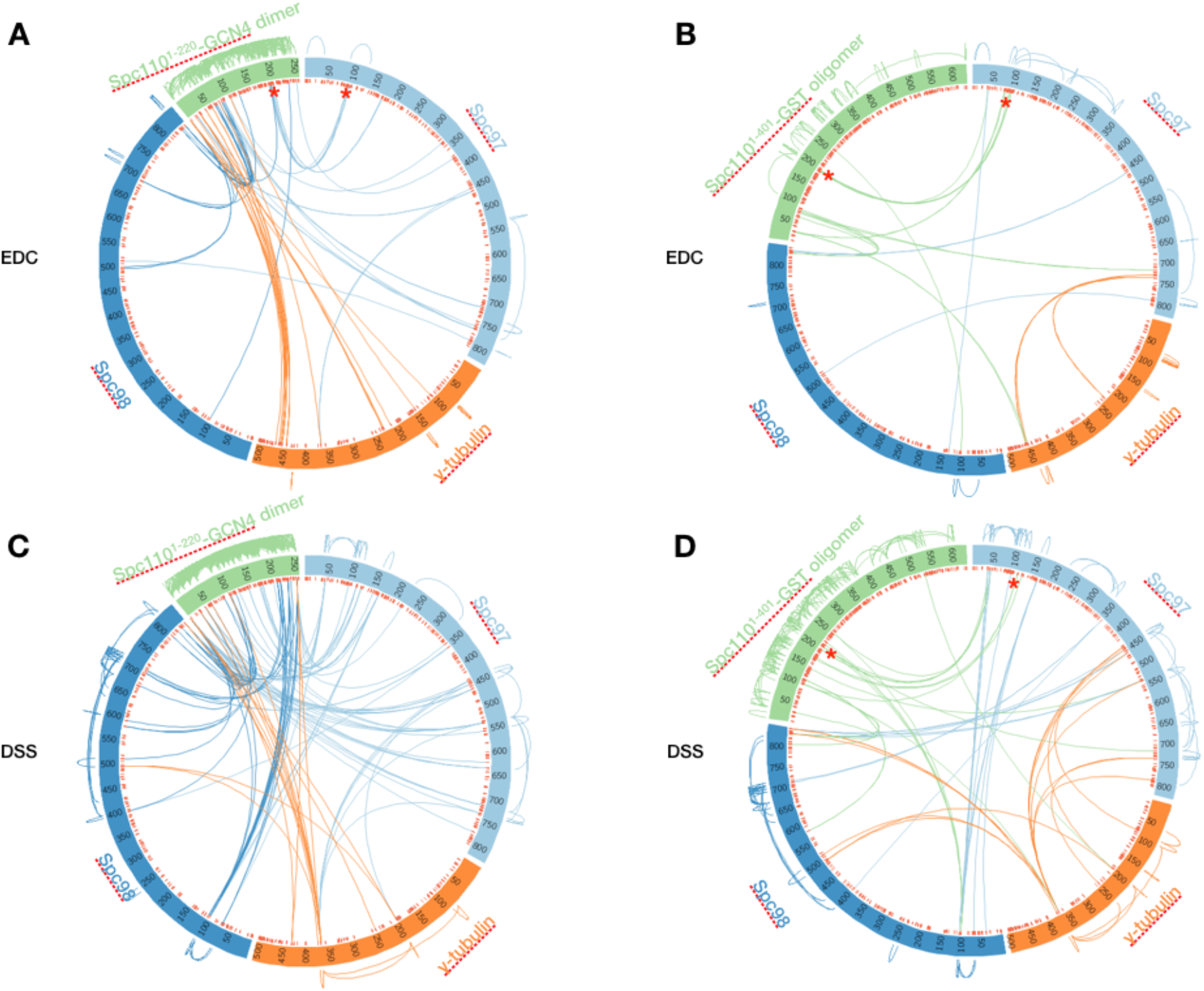
Overview of XL-MS datasets. XL-MS datasets derived from EDC crosslinked (**A-B**) and DSS (**C-D**) crosslinked samples containing γTuSC plus Spc110^1-220^-GCN4 dimer (**A, C**) or Spc110^1-401^-GST (**B, D**). Intermolecular crosslinks are shown within the circle, while intramolecular crosslinks are shown outside the circle. Red ticks inside the circle represent positions that are reactive to the respective crosslinking reagents. The red asterisks show crosslinks between the Spc110 NCC and the N-terminus of Spc97.

### The Spc110 NCC^164-208^ binds to γTuSC at the N-terminal regions of Spc97 and Spc98

Due to the limited resolution of the cryo-EM reconstruction, the derived atomic model contains only the peptide backbone (Kollman, et al., 2015; Greenberg, et al., 2016). Given the proximity between the NCC^164-208^ and γTuSC observed by XL-MS, we sought a higher-resolution structure of the NCC region via X-ray crystallography. Previous work indicates that Spc110^1-220^ is only weakly dimeric (Lyon, et al., 2016). We thus screened several coiled-coil domain fusion constructs of the Spc110 NCC (Figure 3A). These fusion domains have been shown to aid crystallization of weakly interacting coiled-coils (Frye, et al., 2010; Klenchin, et al., 2011). N-terminal fusions with Xrcc4 and Gp7 produced high yields of soluble protein. We elected to move on with the Xrcc4 fusion as it contained a longer portion of the NCC (residues 164-207). The Xrcc4-Spc110^164-207^ construct crystallized in a variety of conditions. Crystals yielded diffraction data to 2.1 Å and phases were obtained by molecular replacement using the Xrcc4 structure as a search model. As expected, the electron density map was consistent with a coiled coil, with interpretable density for Spc110 residues 164-203.

**Figure 3.**
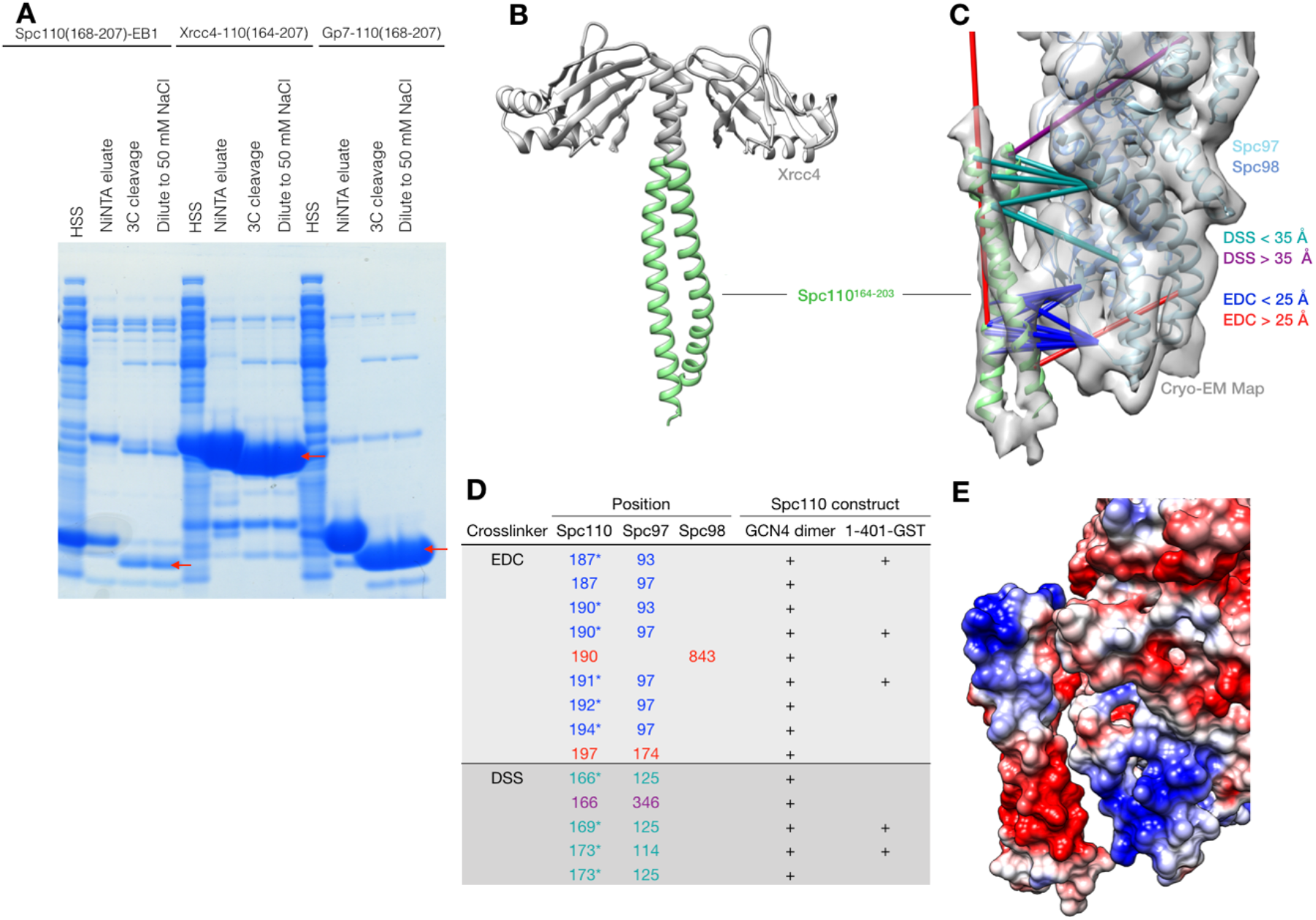
Spc110 NCC structure determination. **A**. Screening Spc110 NCC^164-208^ coiled-coil domain fusion constructs. Constructs were expressed with N-terminal 6His affinity tags in *E. coli* then lysed, centrifuged to produce a high-speed supernatant (HSS), then purified by nickel affinity chromatography. Eluates from nickel affinity columns were incubated with 3C protease to cleave the 6His tag, then diluted with NaCl-free buffer to 50 mM NaCl before further purification. **B**. Structure of Xrcc4-Spc110^164-207^, where Spc110 NCC residues 164-203 are resolved. **C**. Spc110 NCC structure fit into γTuSC cryo-EM density map (grey surface, EMDB ID 2799) along with γTuSC pseudo-atomic model (PDB ID 5FLZ) (Kollman, et al., 2015; Greenberg, et al., 2016). The majority of XL-MS distance restraints are satisfied by this model. Satisfied and violated DSS crosslinks are shown in cyan and purple, respectively. Satisfied and violated EDC crosslinks are shown in blue and red, respectively. Crosslinks that are satisfied by either Spc110 monomer are shown twice, one for each monomer. **D**. Table of crosslinks between Spc110 NCC^164-208^ and γTuSC. Spc110 crosslinks marked with an asterisk (*) are satisfied on either monomer within NCC^164-208^. **E**. Coulombic potential map of Spc110 NCC^164-208^ and γTuSC, showing charge complementarity between acidic patch on Spc110 and basic patch on γTuSC. Map was calculated using UCSF Chimera (Pettersen, et al., 2004).

When docked into the cryo-EM map, the X-ray model occupies most of the alpha-helical cryo-EM density. We then mapped the 14 unique DSS and EDC crosslinks onto the combined γTuSC-Spc110 NCC^164-203^ model. The majority of both DSS (4/5) and EDC (7/9) crosslinks are within expected C_α_-C_α_ distances (<25 Å for EDC, <35 Å for DSS) (Figure 3C). From the crosslinking data and the fact that the NCC is resolved in the cryo-EM map, we conclude that the Spc110 NCC domain forms a stable contact with γTuSC. The Coulombic surface representation of Spc110 NCC and the N-terminal regions of Spc97 and Spc98 of γTuSC suggests the interactions are mediated by charge complementarity between acidic residues in Spc110 and basic residues in γTuSC (Figure 3D).

### Integrative structural model of γTuSC-Spc110 based on cryo-EM, X-ray crystallography, and XL-MS

Given the large number of distance restraints in our XL-MS dataset and the availability of structural models for the Spc110 NCC^164-208^ and γTuSC, we next sought to generate a structural model of the γTuSC-Spc110 complex using integrative modeling (Alber, et al., 2007; Russel, et al., 2012). The γTuSC was represented by the pseudo-atomic structure inferred from a cryo-EM map of the closed, disulfide-stabilized γTuSC filament and remotely related template structures (PDB ID: 5FLZ) (Kollman, et al., 2015; Greenberg, et al., 2016) and the Spc110 NCC^164-208^ was represented by its crystal structure (Figure 3B). Regions with unknown structure were modeled as flexible strings of beads. For Spc110, each sphere represented 5 residues. Finally, the proximity between specific residue pairs was determined by DSS and EDC XL-MS experiments using Spc110^1-401^-GST. We used the Spc110^1-401^-GST crosslinks as there were fewer intra-Spc110 crosslinks than with the Spc110^1-220^-GCN4 dimer (Figure 2). The NCC^164-203^ was docked into the cryo-EM density map by cross-correlation based fitting in Chimera (Pettersen, et al., 2004) and its position was not optimized as part of the integrative modeling process. The position of γTuSC and the NCC^164-203^ was fixed during sampling, while the positions of flexible beads in the regions Spc110^1-163^ and Spc110^204-220^ were optimized. Next, 1.5 million γTuSC-Spc110 models were computed by optimizing spatial proximities, as informed by cross-linking data, excluded volume, and sequence connectivity from 50 random initial models. The top-scoring one thousand models, which sufficiently satisfy the information used to compute the models, were used for analysis. Details of the integrative modeling process are presented in Supplementary Computational Methods.

The ensemble of models clustered predominantly in one class (80.5% of models) with an overall localization precision of 46.4 Å, where the precision is defined as the average bead RMSD between all pairs of models in the cluster. All except 2 of 15 EDC crosslinks between Spc110 and γTuSC are satisfied in the top-scoring model in the cluster; these 2 crosslinks (between Spc110^24^ and Spc97^80^/Spc97^82^) likely represent false-positives or reflect a structural state not accessible by our representation (e.g., due to modeling with rigid body representations of complex components or due to actual assemblies containing more than one γTuSC). Similarly, only 1 out of 17 DSS crosslinks (between Spc97^716^ and the Spc110 NCC^164-208^) is violated. In general, an ensemble of models can be visualized as a localization probability density map. The map specifies the probability of any volume element being occupied by a given bead in superposed good-scoring models. In the localization probability density map for the entire cluster, the position of the Spc110 NTD^1-111^s appears blurred, especially at higher contour levels (Figure 4A). This uncertainty is explained in part by the fact that, in the top scoring-model in the cluster, only one Spc110 NTD^1-111^ is closely associated with γTuSC (Figure 4B, dark green) while the other appears to have an unconstrained localization in the solvent away from γTuSC (Figure 4B, light green). This result shows that most crosslink distance restraints can be satisfied by a single Spc110 NTD, raising questions about how multiple copies of NTD^1-111^ work together to stabilize γTuRC.

**Figure 4.**
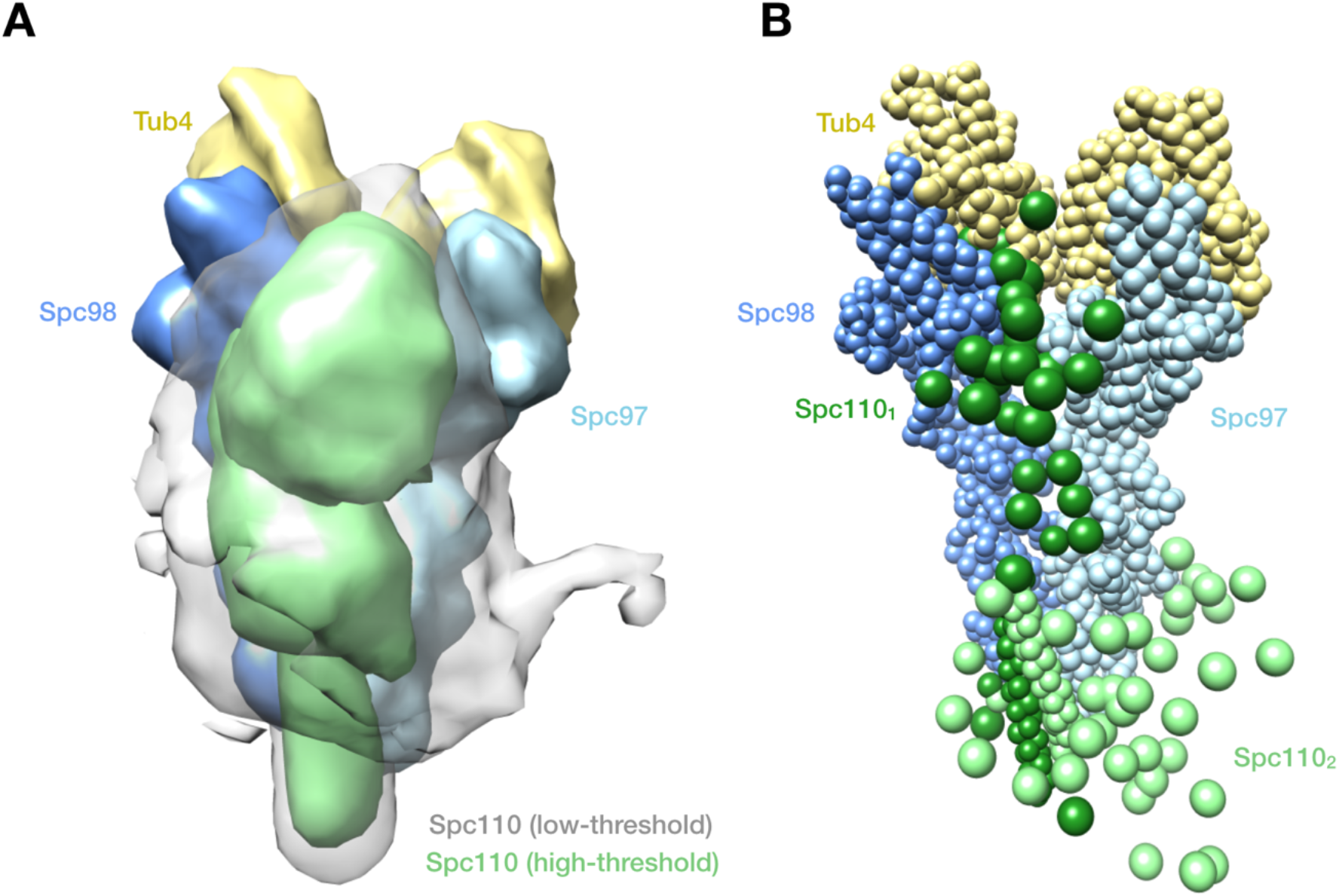
Integrative structural model of γTuSC-Spc110 complex. **A**. Localization density map showing the positions of different parts of the complex in the ensemble of models that sufficiently satisfy the data. Spc110 is shown divided into three regions (residues 1-163, 164-203, and 204-220) and shown at two contour levels. The light green surface shows the localization density contour at 25% of the maximum voxel value, while the transparent grey surface shows the contour at 10% of the maximum voxel value. **B**. Top-scoring model within the most-occupied cluster in bead representation, showing one Spc110 NTD^1-111^ (light green) not associated with γTuSC.

### The role of a conserved cysteine in Spc110

We observed crosslinks between the Spc110 central coiled coil domain and the N-terminal domains of Spc97 and Spc98 within γTuSC (Figure 2B and 2D). These domains of Spc97 and Spc98 are unresolved in the cryo-EM reconstruction (Kollman, et al., 2015). This region of Spc98 is important for Spc110-mediated γTuRC assembly (Lin, et al., 2015). We thus wanted to include this region of Spc110 in further experiments. This required mutating a conserved cysteine at Spc110 position 225 (Figure 5A), as we found that disulfides formed between the cysteines leading to higher-order oligomerization that confounds analysis of other features of Spc110 (Figure 5B-C). C225 is one of two cysteine residues in Spc110, the other being C911 in the calmodulin binding site near the C-terminus, and Spc110 has been shown to form disulfides in vivo (Knop & Schiebel, 1997). C225 is in an intriguing position within Spc110, as there is a predicted break in the coiled-coil near this position (Supplementary Figure S1), suggesting disulfides form not within an Spc110, but between adjacent Spc110 dimers to stabilize higher-order oligomers. However, the Spc110 C225S mutant was viable *in vivo* as assessed by a plasmid shuffle assay, as was a mutation of residues 224 and 226 to lysine, which should generate a hyper-reactive KCK motif, indicating that neither disulfide formation nor wild-type reactivity of C225 plays an essential role in Spc110 function *in vivo* (Figure 5D).

**Figure 5.**
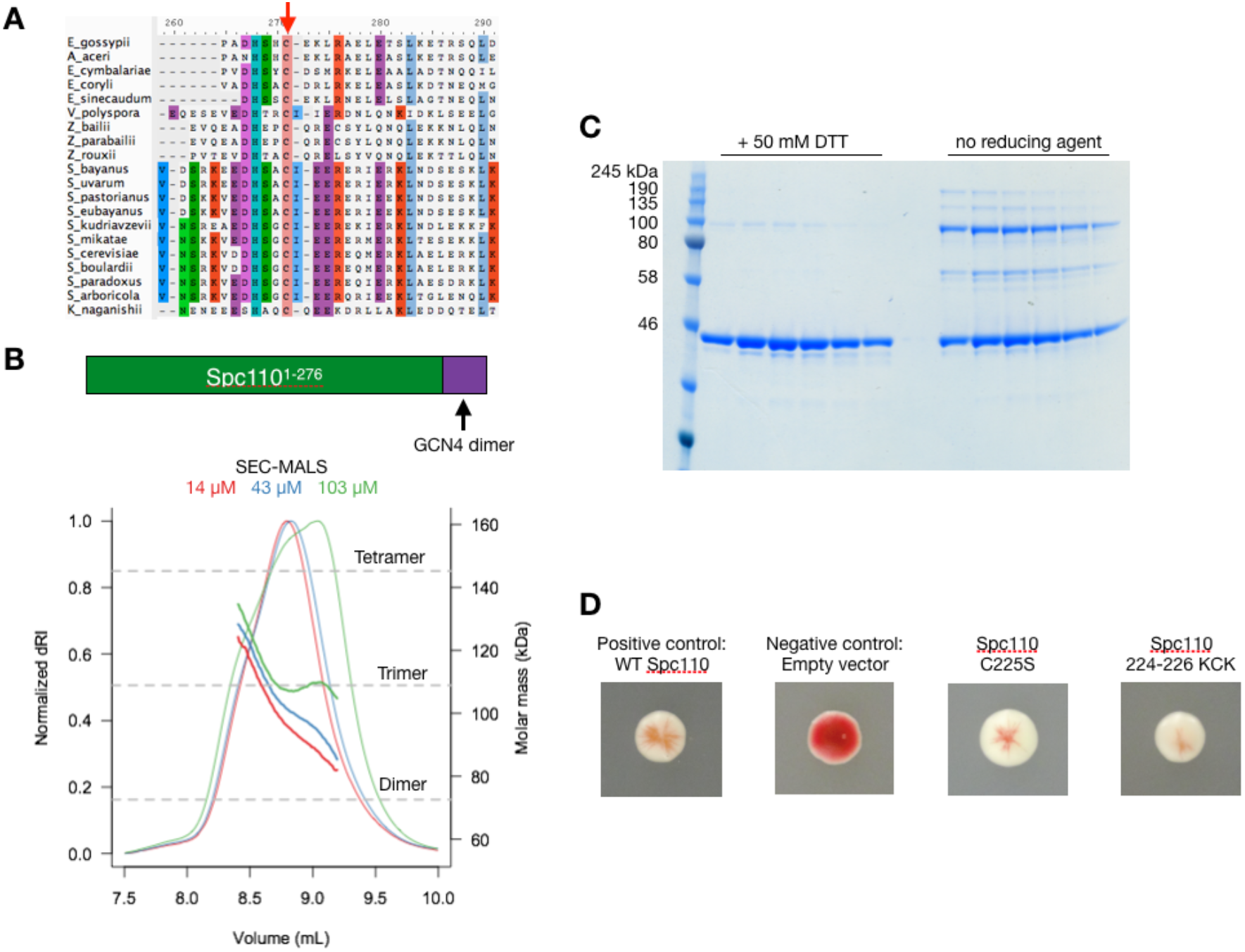
Spc110 contains a non-essential cysteine that forms disulfides in vitro. **A**. Alignment of a variety of fungal Spc110 sequences showing conservation of the cysteine at *S. cerevisiae* position 225. **B**. SEC-MALS analysis of Spc110^1-276^-GCN4 dimer showing formation of species beyond a dimer. **C**. Non-reducing SDS-PAGE shows Spc110^1-276^-GCN4 dimer forms disulfide bonds mediated by cysteine 225, the only cysteine in the construct. **D**. Red-white sectoring plasmid shuffle assay for Spc110 cysteine mutant viability. Strains bearing functional plasmid-encoded Spc110 sector white, indicating both the C225S and the hyper-reactive 224-226 KCK variant are both viable mutants.

### Testing independent effects of the Spc110 NTD^1-111^s

While our integrative structural model does not conclusively prove that only a single Spc110 NTD^1-111^ is required to bind γTuSC, it suggests the possibility that the two NTD^1-111^s within a dimer may serve separate purposes. In particular, the close association of one of the NTD^1-111^s with γTuSC suggests that it acts along with the γTuSC-NCC^164-208^ interaction to stabilize Spc110-γTuSC binding (Figure 4B, dark green). The unassociated NTD^1-111^ could then be free to stabilize interactions between adjacent γTuSCs, potentially explaining the decrease in average γTuSC assembly size observed with the Spc110-Δ111 truncation mutant (Lyon, et al., 2016).

To address these possibilities, we turned to the SpyCatcher-SpyTag system, which has proven useful in understanding asymmetric behavior by homodimeric proteins (Zakeri, et al., 2012; Elnatan, et al., 2017). We generated fusions between Spc110-C225S residues 1-276 and SpyCatcher or SpyTag domains with short, flexible serine/glycine linkers designed to prevent disruption of the coiled-coil domain by geometric mismatch between the domains. SpyCatcher-SpyTag covalent adducts of Spc110^1-276^-C225S formed readily, allowing comparison of full-length/Δ111 heterodimers with the full-length/full-length or Δ111/Δ111 covalent dimers (Figure 6A). Our FRET assay for γTuRC assembly was then used to analyze the impact of these truncations. This assay uses γTuSC with YFP and CFP fused to the C-termini of Spc97 and Spc98, respectively. As the concentration of Spc110 in the reaction increases, γTuSCs assemble into larger complexes, bringing the fluorophores into proximity and causing increased FRET. The assay provides quantitative information about the apparent affinity of Spc110 for γTuSC from the Spc110 concentration at half-maximal FRET, as well as the average assembly size of γTuSC-Spc110 complexes from the maximal FRET signal (Lyon, et al., 2016) (see Figure S2). At 50 nM γTuSC, the full-length/Δ111 heterodimer reduced the apparent affinity for γTuSC assembly as well as the maximum FRET signal compared with the full-length/full-length control (Figure 6C). The reduced maximum FRET signal of the full-length/Δ111 binding curve implies that the average size of γTuSC assemblies is smaller, and thus that wild-type γTuSC assembly size depends on the presence of two Spc110 NTDs. Consistent with our previous results that only a deletion of residues 1-146 completely abolishes γTuRC assembly (Lyon, et al., 2016), the Δ111/Δ111 dimer continued to stabilize γTuRC assembly very weakly. Our interpretation of these results in summarized in a model (Figure 7) discussed below.

**Figure 6.**
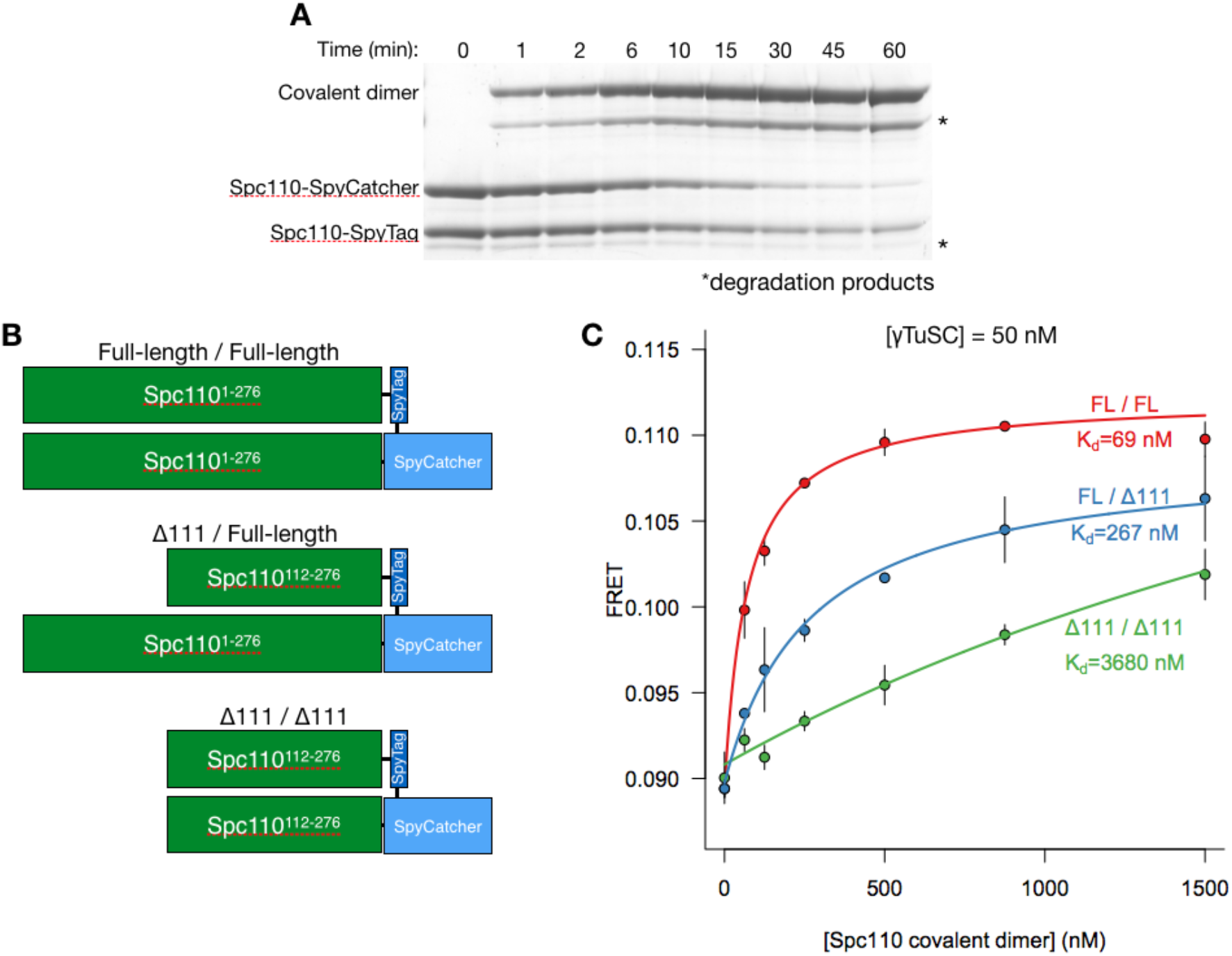
Full-length/Δ111 Spc110 heterodimers show impaired affinity and average assembly size in γTuRC assembly assay. **A**. Time-course of covalent SpyCatcher-SpyTag adduct formation assayed by SDS-PAGE. The asterisk marks a degradation product which is subsequently removed by anion exchange and size exclusion chromatography. **B**. Diagrams of Spc110 covalent dimers. **C**. FRET assay for γTuRC assembly in the presence of Spc110 covalent heterodimers.

**Figure 7.**
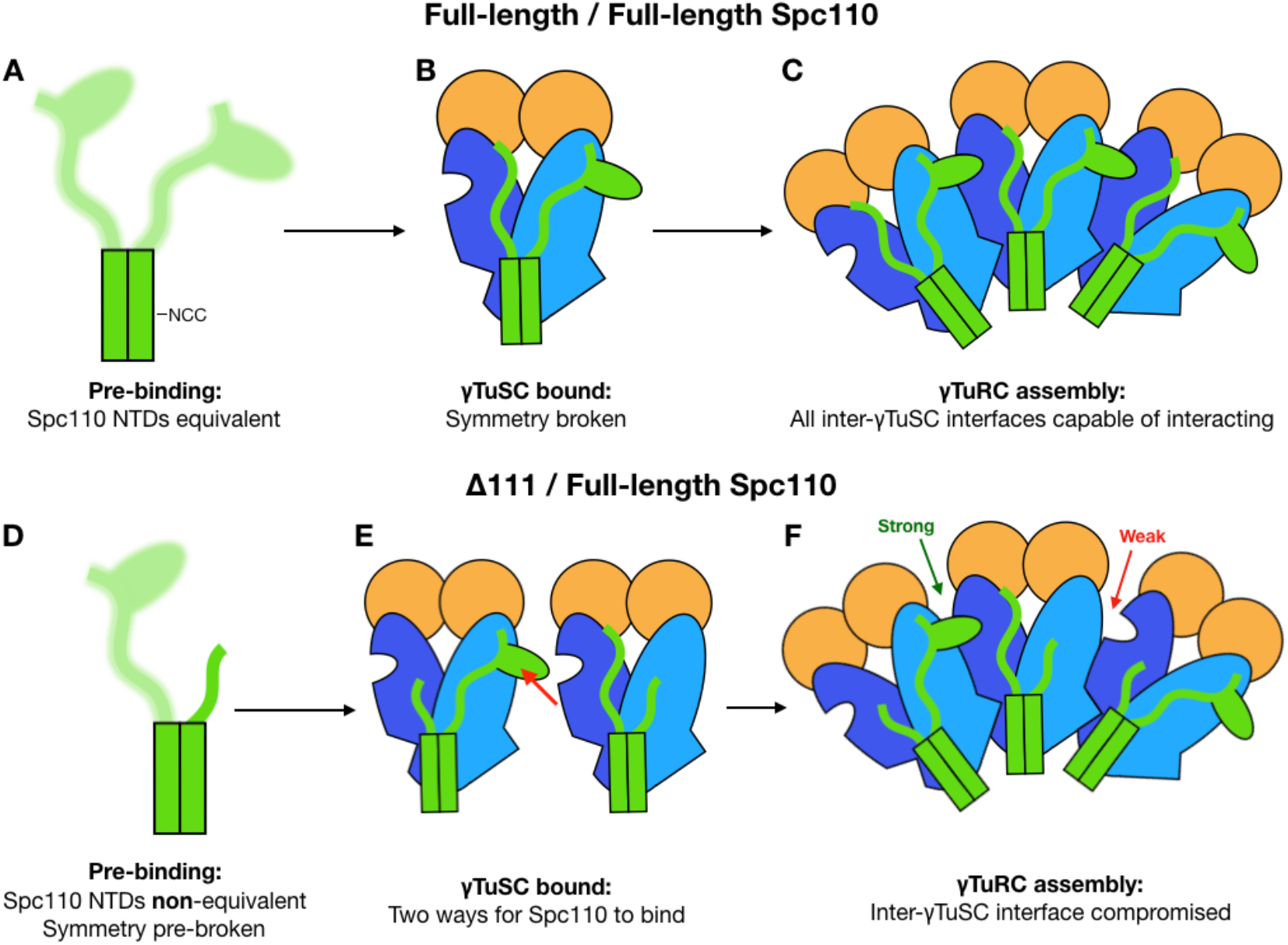
Model for independent action of Spc110 NTD^1-111^s in stabilizing γTuRC. One NTD^1-111^ associates with the same γTuSC which the NCC^164-208^ binds, while the second NTD^1-111^ stabilizes the inter-γTuSC interface. Note that the “bumps-into-holes” representation of Spc110 interacting with γTuSC is purely conceptual in nature and merely represents a stabilizing interaction, not any specific site or mode of interaction.

## Discussion

### Spc110 NTDs serve different roles in γTuRC assembly

The crucial difference between the full-length/full-length dimer and the Δ111/full-length heterodimer is a change in maximum FRET value (Figure 6C). If deleting one NTD^1-111^ simply weakened interaction between Spc110 and γTuSC, the only expected change in the γTuRC assembly curves would be decreased apparent affinity and both Spc110 constructs would reach the same maximum FRET signal at sufficiently high Spc110 concentrations. Importantly, this is not what we observe. As detailed in Supplementary Figure S2, a decrease in maximum FRET implies a decrease in the average number of γTuSCs per Spc110 complex. Our results therefore indicate that the NTD^1-111^ stabilizes γTuSC-γTuSC interactions as stabilizing this interface would lead to an increased average number of γTuSCs per complex at saturating Spc110 concentration.

Based on our results, we propose an independent action model for the two Spc110 NTD^1-111^s (Figure 7). In a full-length/full-length Spc110 dimer, both NTD^1-111^s are equivalent prior to γTuSC binding (Figure 7A). Upon γTuSC binding, this symmetry is broken and one Spc110 NTD makes contacts with the same γTuSC bound by the Spc110 NCC^164-208^, while the other NTD^1-111^ is poised to make contacts with an adjacent γTuSC (Figure 7B). In this scenario, γTuSC self-interactions and the second Spc110 NTD^1-111^ cooperate to stabilize assembly of γTuRC (Figure 7C).

Removing one of the NTD^1-111^s destroys the symmetry that exists in the full-length/full-length dimer (compare Figure 7A with 7D). Thus, Spc110 can bind γTuSC in two configurations (Figure 7E). Only one configuration has an Spc110 NTD^1-111^ poised to contact another γTuSC (Figure 7E, red arrow), leading to γTuRC assemblies where some interfaces are strong due to cooperative stabilization by both γTuSC self-interaction and Spc110, and some are weak due to the absence of additional stabilization by Spc110 (Figure 7F). In the case of the Δ111/Δ111 dimer, γTuRC assembly is severely compromised because only the NCC^164-208^ interaction and the CM1^117-146^ domain are present to interact with γTuSC.

Our XL-MS results and modeling indicate that only one NTD^1-111^ is sufficient to satisfy all the crosslink distance restraints (Figure 4B). Where, then, does the second NTD^1-111^ bind in order to stabilize the inter-γTuSC interface? One possibility is that the observed NTD^1-111^-γTuSC crosslinks in fact represent a mixture of crosslinks from the NTD^1-111^ acting in *cis* and the NTD^1-111^ acting in *trans* from an adjacent γTuSC. Again, the fact that the maximum FRET signal decreases in the Δ111/full-length heterodimer implies that one of the NTD^1-111^s must stabilize the inter-γTuSC interface, which disfavors a model where the NTD^1-111^-γTuSC crosslinks are a mixture of crosslinks from two NTD^1-111^s within a dimer acting in *cis*. The specific nature of the interaction between the NTD^1-111^ and γTuSC requires higher-resolution structural studies.

As an alternative class of models, the NTD^1-111^ may not directly bridge adjacent γTuSCs, but instead enhance the inter-γTuSC self-interaction *via* allostery. If NTD^1-111^ binding allosterically regulates the γTuSC self-interaction interface to enhance its affinity, any perturbation that decreases the residence time of the NTD^1-111^ on γTuSC would be predicted to decrease γTuSC self-interaction affinity. In full-length/full-length Spc110 dimers, when one NTD^1-111^ dissociates there are two NTD^1-111^s which could rebind to γTuSC. However, only one NTD^1-111^ is available in the Δ111/full-length heterodimer, which would lead to reduced residence time on γTuSC, weakened γTuSC self-interaction affinity, and reduced maximum FRET signal. Higher-resolution structural data of γTuSC alone and with Spc110 would aid identification of any such allosteric conformational change mechanisms.

### The Spc110 NCC is a major γTuSC-interacting motif

Our XL-MS dataset revealed the identity of the coiled-coil density previously observed in contact with the N-terminal regions of γTuSC (Figure 3C; Kollman et al. 2015). Unlike the CM1^117-146^ domain, which we previously proposed to be an important γTuSC interaction motif, the NCC^164-208^ contains several mitosis-specific phosphorylation sites (Davis Lab, unpublished data), so it will be important to characterize the role of phosphorylation in γTuSC-NCC^164-208^ binding.

### Implications for γTuRC assembly and the MT cytoskeleton

Our results add another dimension to the role Spc110 plays in γTuRC assembly. In our previous work (Lyon, et al., 2016), we demonstrated that Spc110’ s higher-order oligomerization is the fundamental determinant of its ability to stabilize γTuRC assembly. Our results here represent an elaboration on this core mechanism of γTuRC assembly. We propose that the Spc110 NTD^1-111^s contribute to both high-affinity binding between Spc110 and γTuSC, and to the stabilization of interactions between adjacent γTuSCs. It remains possible that the NTDs are also required to bind additional factors important for MT nucleation, either directly or indirectly. Given recent attention to non-γTuRC-mediated mechanisms of MT nucleation, any cooperation between Spc110, γTuRC, and factors such as Stu2 will be very informative (Roostalu & Surrey, 2017). The trαns-γTuSC interactions that we propose are made by Spc110 could also be important to the mechanisms that lead to γTuRC closure, which we have shown activates the MT nucleating activity of γTuRC (Kollman, et al., 2015). Further study of these questions is necessary and likely to be fruitful.

## Materials and Methods

### Protein expression constructs

Constructs for expression of γTuSC, Spc110^1-220^-GCN4 dimer, and Spc110^1-401^-GST were previously described (Vinh, et al., 2002; Kollman, et al., 2015; Lyon, et al., 2016). Spc110^1-276^-GCN4 dimer was synthesized by GeneArt (Life Technologies). Spc110^1-276^-SpyTag and SpyCatcher fusion constructs were synthesized with the BioXp instrument (SGI-DNA). These constructs were then cloned into pET28 using Gibson Assembly. Point mutations were constructed by site-directed mutagenesis (Zheng, et al., 2004). Truncation mutants were constructed using the Q5 Site-Directed Mutagenesis kit (New England BioLabs).

### Protein Expression and Purification

Purifications for γTuSC, Spc110^1-220^-GCN4 dimer, and Spc110^1-401^-GST were previously described (Vinh, et al., 2002; Kollman, et al., 2015; Lyon, et al., 2016). Spc110^1-276^-GCN4 dimer was purified as for Spc110^1-220^-GCN4 dimer. Spc110^1-276^-SpyCatcher and SpyTag were transformed into BL21(DE3) CodonPlus RIL (Agilent). For each construct, 3 L of culture in Terrific Broth was grown at 30 °C until reaching OD_600_ 0.3-0.4. The temperature was then decreased to 18 °C. Once the culture had reached OD_600_ 0.6-0.8, expression was induced with 0.6 mM IPTG for 16-18 h. Cells were harvested by centrifugation then resuspended in lysis buffer (50 mM potassium phosphate pH 8, 300 mM NaCl, 5 mM EDTA, 1 mM DTT, 0.3% Tween-20, 1x cOmplete protease inhibitor, EDTA-free (Roche)). Cells were lysed by Emulsiflex C3 (Avestin). Lysate was cleared by ultracentrifugation at 40,000 × g for 30 min in a Type 45Ti roto (Beckman-Coulter). Cleared lysate was applied to cOmplete His-Tag purification resin (Roche) and incubated for 1 h at 4 °C with gentle agitation. The column was then washed with 10 CV lysis buffer followed by 10 CV lysis buffer without Tween-20. Spc110 was then eluted with 4 CV of elution buffer (25 mM Tris pH 8.3, 75 mM NaCl, 5 mM EDTA, 1 mM DTT, 1x cOmplete protease inhibitor, EDTA-free (Roche), and 250 mM imidazole). Eluates were then diluted to < 5 mS/cm conductivity with MonoQ buffer A (25 mM Tris pH 8.3, 1 mM DTT). The diluted eluates were then applied separately to a MonoQ 10/100 GL preequilibrated in 2.5% MonoQ buffer B (25 mM Tris pH 8.3, 1 M NaCl, 1 mM DTT) in MonoQ buffer A. The column was then washed with 2 CV of 2.5% MonoQ buffer B, then eluted with a linear gradient from 2.5-50% MonoQ buffer B. Spc110^1-276^ SpyCatcher and SpyTag typically elute at approximately 17 mS/cm and 9 mS/cm conductivity, respectively. The concentration of the pooled fractions containing Spc110^1-276^-SpyCatcher or SpyTag were measured by using Bradford protein assay reagent (Bio-Rad), then combined in a 1:1 molar ratio with the addition of TEV protease to cleave the His-tags. After 1 h, the Spc110 covalent adduct was further purified by size exclusion chromatography on S200 16/60 pg equilibrated in HB150 + 10% glycerol. Fractions containing undegraded Spc110 covalent adducts were then pooled, centrifugally concentrated, flash frozen in liquid nitrogen, and stored at −80 °C. Spc110 coiled-coil “stopper” constructs were purified in essentially the same manner via NiNTA affinity, anion exchange, and size exclusion chromatography.

### Cross-linking and mass spectrometry (XL-MS)

XL-MS was carried out as described by (Zelter, et al., 2015). All γTuSC-Spc110 reactions were in 40 mM HEPES pH 7.0, 150 mM KCl and contained a final concentration 0.4 uM γTuSC and 0.8 uM Spc110. DSS reactions were carried out at room temperature (RT) for 3 min using 0.44 mM DSS prior to quenching with 100 mM ammonium bicarbonate. EDC reactions were carried out at RT for 30 min using 5.4 mM EDC plus 2.7 mM Sulfo-NHS prior to quenching with 100 mM ammonium bicarbonate plus 20 mM 2-mercaptoethanol. After quenching, reactions were reduced for 30 min at 37 °C with 10 mM dithiothreitol (DTT) and alkylated for 30 min at RT with 15 mM iodoacetamide. Trypsin digestion was performed at 37°C for 4 or 6 hours with shaking at a substrate to enzyme ratio of 17:1 or 30:1 for EDC and DSS reactions, respectively, prior to acidification with 5 M HCl. Digested samples were stored at −80°C until analysis. Mass spectrometry and data analysis were performed as described by (Zelter, et al., 2015). In brief 0.25 μg of sample was loaded onto a fused-silica capillary tip column (75-μm i.d.) packed with 30 cm of Reprosil-Pur C18-AQ (3-μm bead diameter, Dr. Maisch) and eluted at 0.25 μL/min using an acetonitrile gradient. Mass spectrometry was performed on a QExactive HF (Thermo Fisher Scientific) in data dependent mode and spectra converted to mzML using msconvert from ProteoWizard (Chambers, et al., 2012).

Proteins present in the sample were identified using Comet (Eng, et al., 2012). Cross-linked peptides were identified within those proteins using Kojak versions 1.42 or 1.4.3 (Hoopmann, et al., 2015) available at http://www.kojak-ms.org. Percolator version 2.08 (Käll, et al., 2007) was used to assign a statistically meaningful *q* value to each peptide spectrum match (PSM) through analysis of the target and decoy PSM distributions. Target databases consisted of all proteins identified in the sample analyzed. Decoy databases consisted of the corresponding set of reversed protein sequences. Data were filtered to show hits to the target proteins that had a Percolator assigned peptide level *q* value ≤ 0.01 and a minimum of 2 PSMs. The complete list of all PSMs and their Percolator assigned q values are available on the ProXL web application (Riffle, et al., 2016) at https://proxl.yeastrc.org/proxl/viewProject.do?project_id=63 along with the raw MS spectra and search parameters used.

### Crystallography

Crystals of Xrcc4-Spc110^164-207^ were obtained with by hanging drop vapor diffusion with 8 mg/mL protein and a well solution containing 13% PEG3350 and 0.2 M magnesium formate. Crystals were cryo-protected by rapid transfer to well solution with 30% PEG3350. Diffraction data was collected under cryogenic conditions at Advanced Light Source beamline 8.3.1. Diffraction data was processed with XDS (Kabsch, 2010) and indexed in space group P1. Phases were obtained by molecular replacement using Phaser within the Phenix package (Adams, et al., 2010; McCoy, et al., 2007). The search model was the PDB ID 1FU1 residues 1-150, with the coiled-coil residues 133-150 mutated to alanine. The S-(dimethylarsenic)cysteine at position 130 in 1FU1 was modified to cysteine. The majority of the structure was built with phenix.autobuild (Terwilliger, et al., 2008) with the remainder built manually in Coot (Emsley, et al., 2010) and refined with phenix.refine (Afonine, et al., 2012). The final structure contains Spc110 residues 164-203, along with the Xrcc4 fusion domain.

### Integrative structural modeling

Integrative structure modeling is described in detail in Supplementary Computational Methods.

### SEC-MALS

SEC-MALS was performed as described using a Shodex Protein KW-804 column and DAWN HELEOS II and OptiLab t-Rex instruments (Wyatt Technology) (Lyon, et al., 2016). The mobile phase was HB150 with 1 mM DTT.

### Sequence alignments

Spc110 sequences were obtained by reciprocal best BLAST (Camacho, et al., 2009) searches with *S. cerevisiae* Spc110 protein sequence. Sequences were aligned using MAFFT version 7.222 (Katoh & Standley, 2013).

### Red-White Sectoring Plasmid Shuffle Assay

Viability of Spc110 mutants was performed with a red-white colony sectoring assay as described (Tien, et al., 2013; Lyon, et al., 2016).

## Author Contributions

ASL and AM created expression constructs, purified proteins, and performed biochemical analyses. AZ performed protein crosslinking and mass spectrometry with RJ. SV performed integrative modeling. KCBY performed the red/white plasmid shuffle assay. ASL, SV, and AZ wrote the paper with input from EM, TND, DAA, and AS. TND, EM, MM, AS, and DAA supervised research.

## Acknowledgements

We gratefully acknowledge many helpful discussions with members of the Agard and Davis labs, as well as our collaborators on the Yeast Centrosome – Structure, Assembly, and Function program project grant in the labs of Chip Asbury, Ivan Rayment, Andrej Sali, Mark Winey, and Sue Jaspersen. We acknowledge financial support from the following sources: Howard Hughes Medical Institute (David A. Agard), National Institute of General Medical Sciences (NIGMS) R01 GM031627 and R35GM118099 (David A. Agard), NIGMS P41 GM103533 (Trisha Davis), NIGMS R01 GM083960 and P41 GM109824 (Andrej Sali), NIGMS P01 GM105537 (David A. Agard, Trisha Davis, and Andrej Sali), National Science Foundation Graduate Research Fellowship Grant No. 1144247 (Andrew S. Lyon), and UCSF Discovery Fellowship (Andrew S. Lyon).

## Supplementary Information

**Figure S1.**
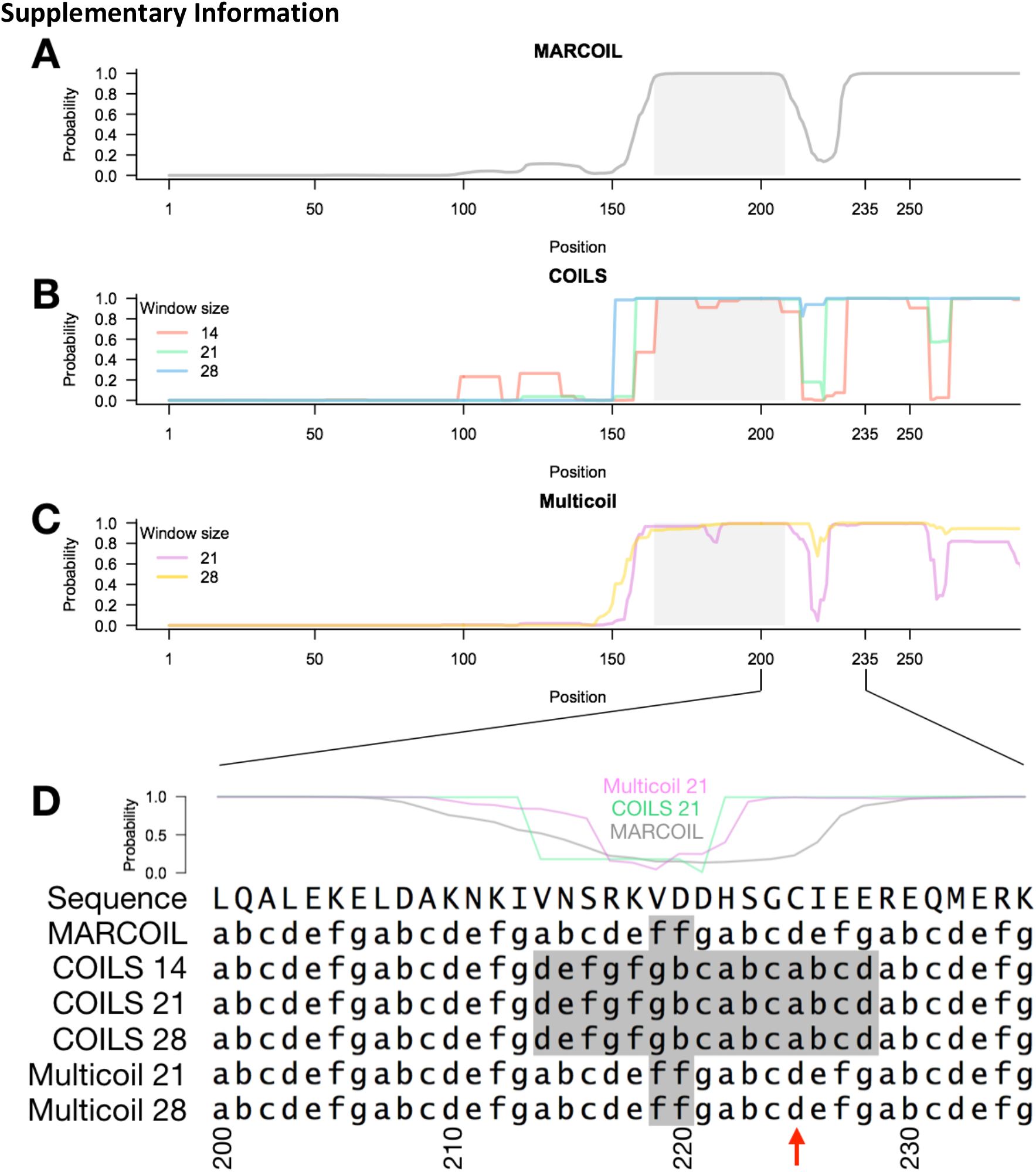
Evidence of discontinuity in the Spc110 coiled coil. **A-C**. Coiled-coil probabilities predicted by MARCOIL, COILS, and Multicoil. All three programs predict a discontinuity around position 220. **D**. Coiled-coil register predicted by MARCOIL, COILS, and Multicoil, with coiled-coil probability plotted above for reference. All three programs predict an interruption in the heptad register around position 220 (grey highlighted positions). Note that C225 (red arrow) is predicted to occupy heptad position a or d, the core positions at the interface between two alpha-helices in a coiled coil. This is inconsistent with the fact that C225S is capable of forming disulfide between Spc110 dimers (Figure 5), further suggesting this region contains a discontinuity in the coiled coil.

**Figure S2.**
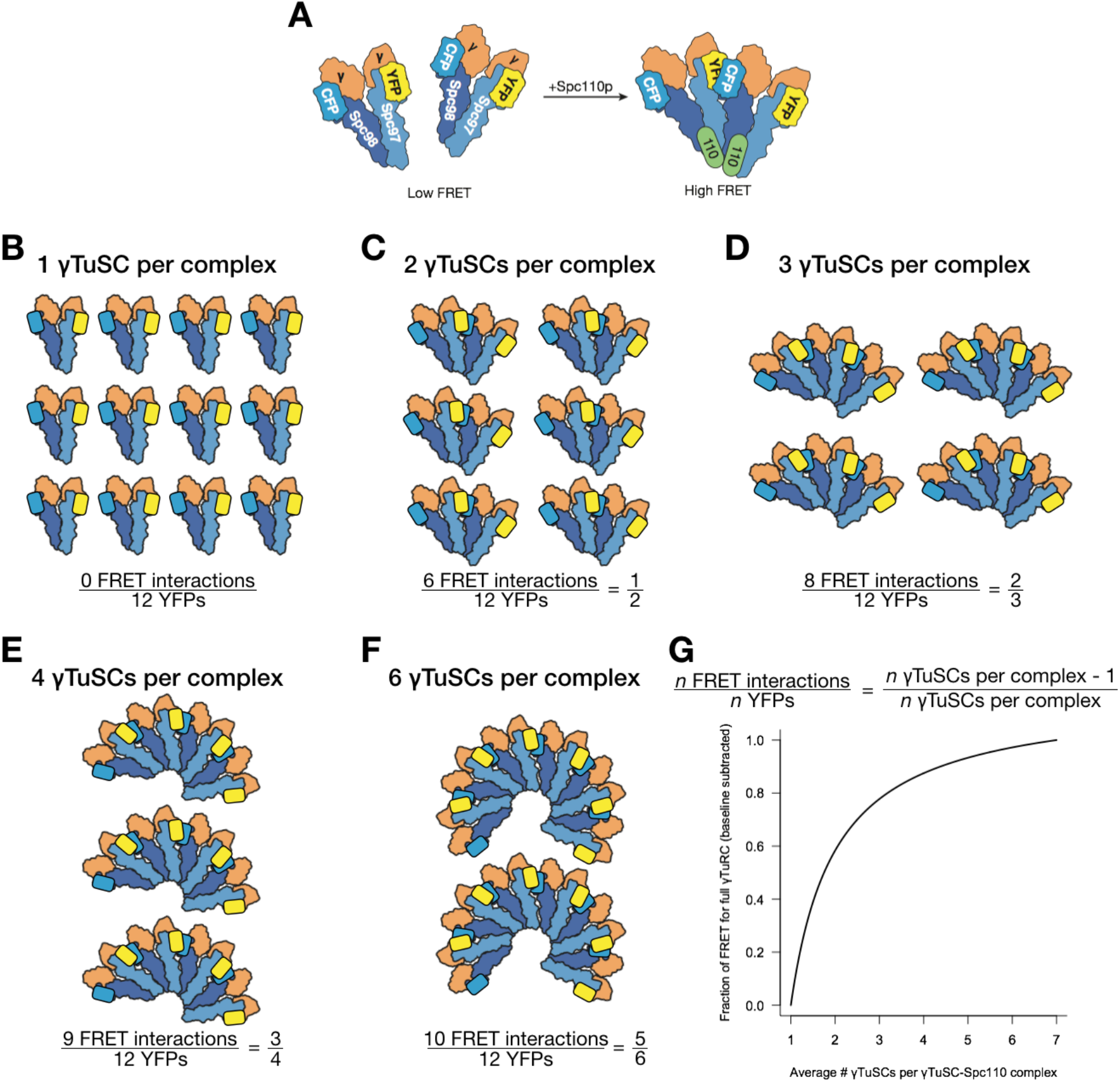
Illustration of FRET assay for γTuRC assembly. **A**. As Spc110 concentration increases, assemblies of γTuSC are stabilized and FRET increases. **B-G**. Demonstration of increases in maximum FRET signal as number of γTuSCs per Spc110 complex increases in a system composed of 12 γTuSCs. For visual clarity, Spc110 is not shown in the cartoons, but in assembled states (**C-F**) γTuSC would be bound by Spc110. **B**. In the unassembled state (one γTuSC per complex), the FRET signal is minimal, with only baseline signal from intra-γTuSC FRET. **C**. When there are two γTuSCs per γTuSC-Spc110 complex, half of YFPs are engaged in an interaction yielding FRET. **D**. With three γTuSCs per γTuSC-Spc110 complex, 2 out of 3 YFPs are engaged in a FRET interaction. The fraction of YFPs engaged in a FRET interaction increases according to a hyperbolic function, with four γTuSCs per complex yielding 3/4 FRET interactions per YFP (**E**) and six γTuSCs per complex yielding 5/6 FRET interactions per YPF (**F**). **G**. Assuming the FRET signal increment for each additional CFP-YFP interaction is independent of γTuSC-Spc110 complex size, and disregarding baseline signal from intra-γTuSC FRET, the relative FRET signal as a function of average γTuSC number (n_γTuSC_) per γTuSC-Spc110 complex is described by the function (n_γTuSC_ – 1)/n_γTuSC_. This function is plotted as a function of n_γTuSC_ and is scaled so that a complete γTuRC (n_γTuSC_ = 7) has a FRET signal of 1.

## Supplementary Computational Methods

### Localizing Spc110 on γTuSC using integrative structure determination

The localization of Spc110 on γTuSC using integrative structure determination proceeded through four stages (Alber *et al*., 2007; Russel *et al*., 2012): (1) gathering data, (2) representing subunits and translating data into spatial restraints, (3) conformational sampling to produce an ensemble of structures that satisfies the restraints, and (4) analyzing and validating the ensemble structures and data. The modeling protocol (*i.e*., stages 2, 3, and 4) was scripted using the *Python Modeling Interface* (PMI) package, a library for modeling macromolecular complexes based on our open-source *Integrative Modeling Platform* (IMP) package, version 2.8 (https://integrativemodeling.org) (Russel *et al*., 2012). The current procedure is an updated version of previously described protocols (Viswanath *et al*., 2017; Wang *et al*., 2017; Kim *et al*., 2018; Webb *et al*., 2018). Files containing the input data, scripts, and output results will be available at https://salilab.org/gtuscSpc110 as well as the nascent Protein Data Bank archive for integrative structures (https://pdb-dev.wwpdb.org/).

#### Stage 1: Gathering data

17 DSS and 15 EDC cross-links (Figure 2) between Spc110 and γTuSC were identified *via* mass spectrometry of Spc110^1-401^-GST, informing the localization of Spc110 relative to γTuSC. γTuSC structure used was obtained from the PDB (code 5FLZ); it was determined primarily based on a cryo-EM density map at 6.9 Å resolution (EMDB code: 2799) (Kollman *et al*., 2015; Greenberg *et al*., 2016). Representation of Spc110 relied on (i) crystal structure of Spc110 N-terminal coiled-coil (NCC) domain (Figure 3) and (ii) failure to detect related sequences of known structure in the rest of the Spc110 sequence by HHPred (Söding *et al*., 2005). Validation of the models relied on biochemical analysis γTuRC assembly with covalent Spc110 heterodimers using the FRET assay (Figure 6).

#### Stage 2: Representing subunits and translating data into spatial restraints

Information about the modeled system (above) can in general be used for defining its representation, defining the scoring function that guides sampling of alternative models, limiting sampling, filtering of good-scoring models obtained by sampling, and final validation of the models. Here, the “flexible” representation for most of Spc110 reflects the absence of known related structures. The γTuSC and Spc110 NCC domain representations rely on their atomic structures. The scoring function relies on chemical cross-links, excluded volume, and sequence connectivity. The validation relies on biochemical analysis and FRET assays for γTuRC assembly.

An optimal representation facilitates accurate formulation of spatial restraints as well as efficient and complete sampling of good-scoring solutions, while retaining sufficient detail without overfitting, so that the resulting models are maximally useful for subsequent biological analysis. To maximize computational efficiency while avoiding using too coarse a representation, we represented the system in a multi-scale fashion. A rigid body consisting of multiple beads was defined for γTuSC and the Spc110 NCC (Spc110^164-203^). In a rigid body, the beads have their relative distances constrained during conformational sampling, whereas in a flexible string the beads are restrained by the scoring function (below). Rigid bodies were coarse-grained using one-residue beads, whose coordinates were those of the corresponding C_α_ atoms. The remaining regions in γTuSC without an atomic model were represented by a flexible string of beads encompassing 20 residues each. Due to lack of acceptable comparative models, and knowing that a large region of the N-terminus of Spc110 lacks secondary structure (Figure 1), we used a flexible string of 5-residue beads each to represent regions of Spc110 other than the coiled-coil domains. With this representation in hand, we next encoded the spatial restraints into a scoring function based on the information gathered in Stage 1, as follows:

1. *Cross-link restraints:* The 17 DSS and 15 EDC cross-links (Figure 2) were used to construct the Bayesian scoring function (Rieping *et al*., 2005) that restrained the distances spanned by the cross-linked residues (Shi *et al*., 2014).
2. *Excluded volume restraints:* The excluded volume restraints were applied to each bead, using the statistical relationship between the volume and the number of residues that it covered (Alber *et al*., 2007).
3. *Sequence connectivity restraints:* We applied the sequence connectivity restraints, using a harmonic upper distance bound on the distance between consecutive beads in a subunit, with a threshold distance equal to twice the sum of the radii of the two connected beads. The bead radius was calculated from the excluded volume of the corresponding bead, assuming standard protein density (Alber *et al*., 2007; Shi *et al*., 2014).

#### Stage 3: Conformational sampling to produce an ensemble of structures that satisfies the restraints

We aimed to maximize the precision at which the sampling of good scoring solutions was exhaustive (Stage 4). We sampled the positions of flexible Spc110 beads and the flexible linkers of γTuSC. The search for good-scoring models relied on Gibbs sampling, based on the Metropolis Monte Carlo algorithm (Wang *et al*., 2017). The positions of the γTuSC rigid body and the Spc110 NCC rigid body was fixed, while the initial positions of flexible γTuSC and Spc110 beads were randomized. The Monte Carlo moves included random translation of individual beads in the flexible segments of γTuSC and Spc110 (up to 3 Å). A model was saved every 10 Gibbs sampling steps, each consisting of a cycle of Monte Carlo steps that moved every moving bead once.

This sampling produced a total of 1,500,000 models from 50 independent runs, requiring ~2 days on 200 CPU cores. For the most detailed specification of the sampling procedure, see the IMP modeling script (https://salilab.org/gtuscSpc110). We only consider for further analysis the top-scoring 1,000 models based on total score. This threshold implies satisfaction of the input datasets within their uncertainties (below).

#### Stage 4: Analyzing and validating the ensemble structures and data

Input information and output structures need to be analyzed to estimate structure precision and accuracy, detect inconsistent and missing information, and to suggest more informative future experiments. We used the analysis and validation protocol published earlier (Alber *et al*., 2007; Viswanath *et al*., 2017; Kim *et al*., 2018): Assessment began with the clustering of the models and estimating their precision based on the variability in the ensemble of good-scoring structures, and quantification of the structure fit to the input information. These validations are based on the nascent wwPDB effort on archival, validation, and dissemination of integrative structure models (Sali *et al*., 2015; Burley *et al*., 2017). We now discuss each one of these points in turn.

##### (1) Clustering and structure precision

An ensemble of good-scoring structures needs to be analyzed in terms of the precision of its structural features (Alber *et al*., 2007; Kim *et al*., 2018). The precision of a component position can be quantified by its variation in an ensemble of superposed good-scoring structures. It can also be visualized by the localization probability density for each of the components of the model.

As described above, integrative structure determination of the Spc110-γTuSC complex resulted in effectively a single good-scoring solution, at the precision of 46.4 Å, where the precision is defined by the average bead RMSD between all pairs of models in the cluster (Shi *et al*., 2014).

##### (2) Fit to input information

An accurate structure needs to satisfy the input information used to compute it. First, the top-scoring model of the dominant cluster satisfied 86.67% (94%) of the EDC (DSS) cross-links; an EDC (DSS) cross-link restraint is satisfied by a model if the corresponding Cα-Cα distance in the model (considering restraint ambiguity) is < 25 Å (35 Å). The remainder of the restraints are harmonic, with a specified standard deviation. The cluster generally satisfied at least 95% of restraints of each type (excluded volume and sequence connectivity). A restraint is satisfied by a cluster of models if the restrained distance in any model in the cluster (considering restraint ambiguity) is violated by less than 3 standard deviations, specified for the restraint. Most of the violations are small, and can be rationalized by local structural fluctuations, coarse-grained representation of the model, and/or finite structural sampling.

##### (4) Satisfaction of data and considerations that were not used to compute the models

The most direct test of a modeled structure is by comparing it to the data that were not used to compute it (a generalization of cross-validation). In this regard, model validation was based on the FRET assay for γTuRC assembly with covalent Spc110 heterodimers, which provide support to the conclusion from the model that the two NTDs of a Spc110 dimer act independently (Figure 6).

